# SAFE Acoustics: an open-source, real-time eco-acoustic monitoring network in the tropical rainforests of Borneo

**DOI:** 10.1101/2020.02.27.968867

**Authors:** Sarab S. Sethi, Robert M. Ewers, Nick S. Jones, Aaron Signorelli, Lorenzo Picinali, C. David L. Orme

## Abstract

1. Automated monitoring approaches offer an avenue to deep, large-scale insight into how ecosystems respond to human pressures. Since sensor technology and data analyses are often treated independantly, there are no open-source examples of end-to-end, real-time ecological monitoring networks.
2. Here, we present the complete implementation of an autonomous acoustic monitoring network deployed in the tropical rainforests of Borneo. Real-time audio is uploaded remotely from the field, indexed by a central database, and delivered via an API to a public-facing website.
3. We provide the open-source code and design of our monitoring devices, the central web2py database and the ReactJS website. Furthermore, we demonstrate an extension of this infrastructure to deliver real-time analyses of the eco-acoustic data.
4. By detailing a fully functional, open-source, and extensively tested design, our work will accelerate the rate at which fully autonomous monitoring networks mature from technological curiosities, and towards genuinely impactful tools in ecology.

## Introduction

Natural ecosystems around the world are undergoing rapid changes due to increasing human pressures^1,2^. Monitoring these changes in real time is essential for deploying interventions where and when they are most urgently needed, and for furthering our understanding of complex ecological responses to habitat quality degradation^3^.

Whilst traditional survey methods tend to be slow and laborious^4^, recording and analysing the sounds of an environment – its soundscape – has shown great promise as a tractable route to fully automated and scalable ecological monitoring^5–7^. Here we detail the design and implementation of SAFE Acoustics, a fully automated, real-time, acoustic monitoring network. The network is deployed across a tropical rainforest fragmentation experiment in Sabah, Malaysia^8^, and will be used to probe the effects of logging intensity and agricultural land conversion on biodiversity and ecosystem stability.

The SAFE Acoustics monitoring network can be broadly split into three key components: (i) an array of real-time acoustic monitoring devices deployed in the field (Fig. 1a); (ii) a web server that indexes the incoming audio and provides a standardised machine-readable interface to the data (Fig. 1b), and; (iii) a website which allows the public to browse and listen to the latest audio from the network (Fig. 1c). Additionally, we demonstrate how this infrastructure can be extended to deliver real-time estimates of habitat quality using a fully automated analysis pipeline (Fig. 1d).

**Figure 1:**
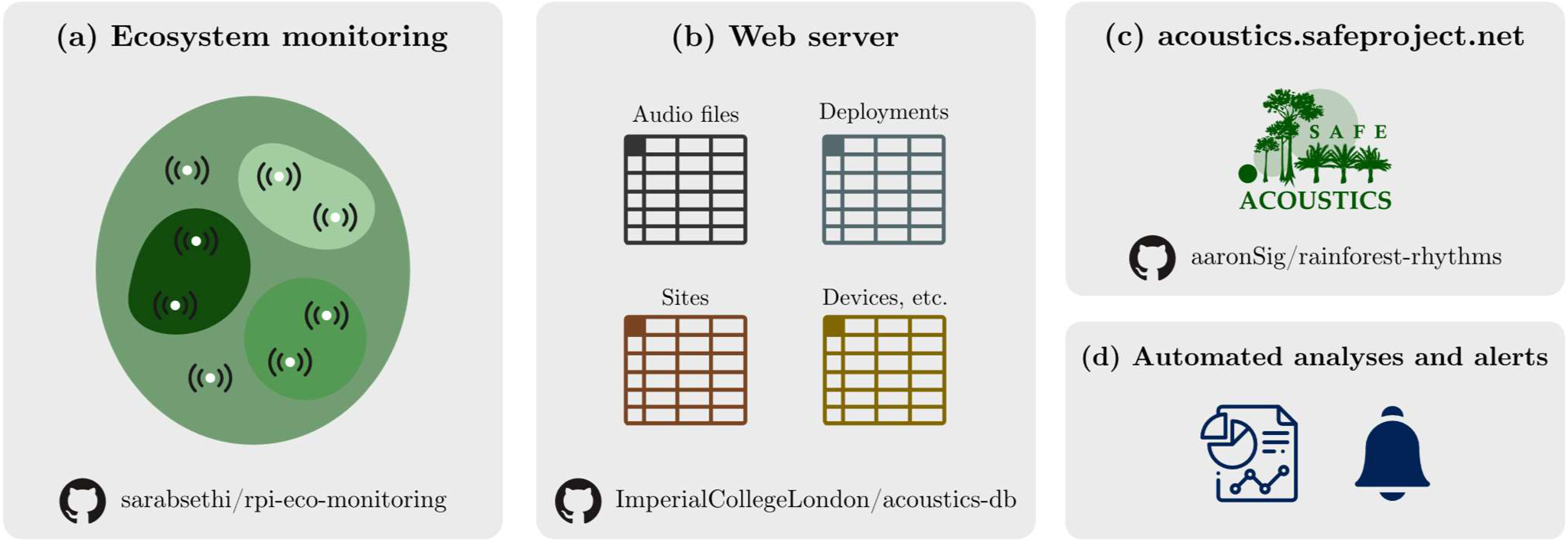
A pipeline for a continuous, real-time acoustic monitoring network. (a) Real-time acoustic monitoring devices record audio from the field and upload files to a remote FTP server. **(b)** A web server continuously scans and indexes the incoming audio and provides a standardised interface to access it. This interface is used by **(c)** a website, that allows the public to browse and listen to the latest audio data, and **(d)** real-time analyses which automatically extract ecological information from the soundscapes. Paths to Github repositories are provided for **(a), (b)**, and **(c)** at the bottom of their respective panels.

Every ecological monitoring effort has its own unique challenges and goals, and developing a one-size-fits-all solution is not possible^9^. Therefore, we have open sourced all the technology developed during our study, including the hardware design and firmware, and complete source code for the web server and public-facing website. Our system provides a fully functional template for a robust and reliable autonomous ecological monitoring network, which others can adjust and build upon to fit the particular needs of their own projects.

### Real-time monitoring devices and server infrastructure

The SAFE Project is a forest fragmentation experiment in Malaysian Borneo, with study sites across old growth, selectively logged, and salvage logged forest, and within nearby palm oil plantations. We deployed eleven Raspberry Pi based autonomous acoustic monitoring devices^7^ across this gradient in February 2018. Each solar-powered device continuously transmits compressed audio files to a remote FTP server, using a 3G mobile internet connection. Data is uploaded within a folder structure specifying the unique device ID, date, and time of each recording. The monitoring devices used are fully open-source, and further information can be found at http://rpi-eco-monitoring.com.

To enable efficient searching and retrieval of the acoustic data, a web2py-based^10^ web app performs a daily scan of the FTP server, and automatically indexes new audio files in an SQL database. An “Audio” table contains an entry for each recording, along with its date, time and the unique device ID it was uploaded from. A “Deployments” table links device IDs to geographical monitoring sites, and a “Sites” table contains metadata such as habitat quality for each location. The web app is also able to index audio from semi-autonomous recording devices (e.g., AudioMoths^11^) using the same database structure. Within the same database we integrate avifaunal and herpetofaunal point-count datasets^12^ to provide a list of expected species at each site. A full inventory of animals within this dataset is stored in a “Taxa” table, and associated metadata and media is linked from the Global Biodiversity Information Facility (GBIF). Full source code for the web app, and detailed documentation is available at https://github.com/ImperialCollegeLondon/acoustics-db.

The web app also provides an application programming interface (acoustics-db API) to provide access to the audio data in a machine-readable format. This allows external applications to retrieve data from the monitoring network in a standardised manner, without needing direct access to the raw data or any prior knowledge of back-end implementation details. The API forms the backbone of both the public-facing website, and the real-time analysis pipeline presented in the following sections. The SAFE Project currently has this server deployed at https://acoustics-db.safeproject.net.

### Public-facing SAFE Acoustics website

To expose the audio collected from our acoustic monitoring network in a user-friendly interface we developed a public-facing website (http://acoustics.safeproject.net/, Fig. 2). The website is implemented using the ReactJS framework and is rendered entirely on the client side to reduce server-side load and associated costs. Direct links to audio files, and the associated metadata is acquired from the acoustic monitoring network through the acoustics-db API. Users of the website are able to browse a map of the recorder locations, listen to audio across the different hours of the day, and explore the species communities at each site. Exploring qualitatively how the soundscape changes across different levels of landscape degradation, and throughout the stages of the diurnal cycle serves as an impactful scientific communication tool. Full implementation details of the website can be found together with the open source code at https://github.com/aaronSig/rainforest-rhythms.

**Figure 2:**
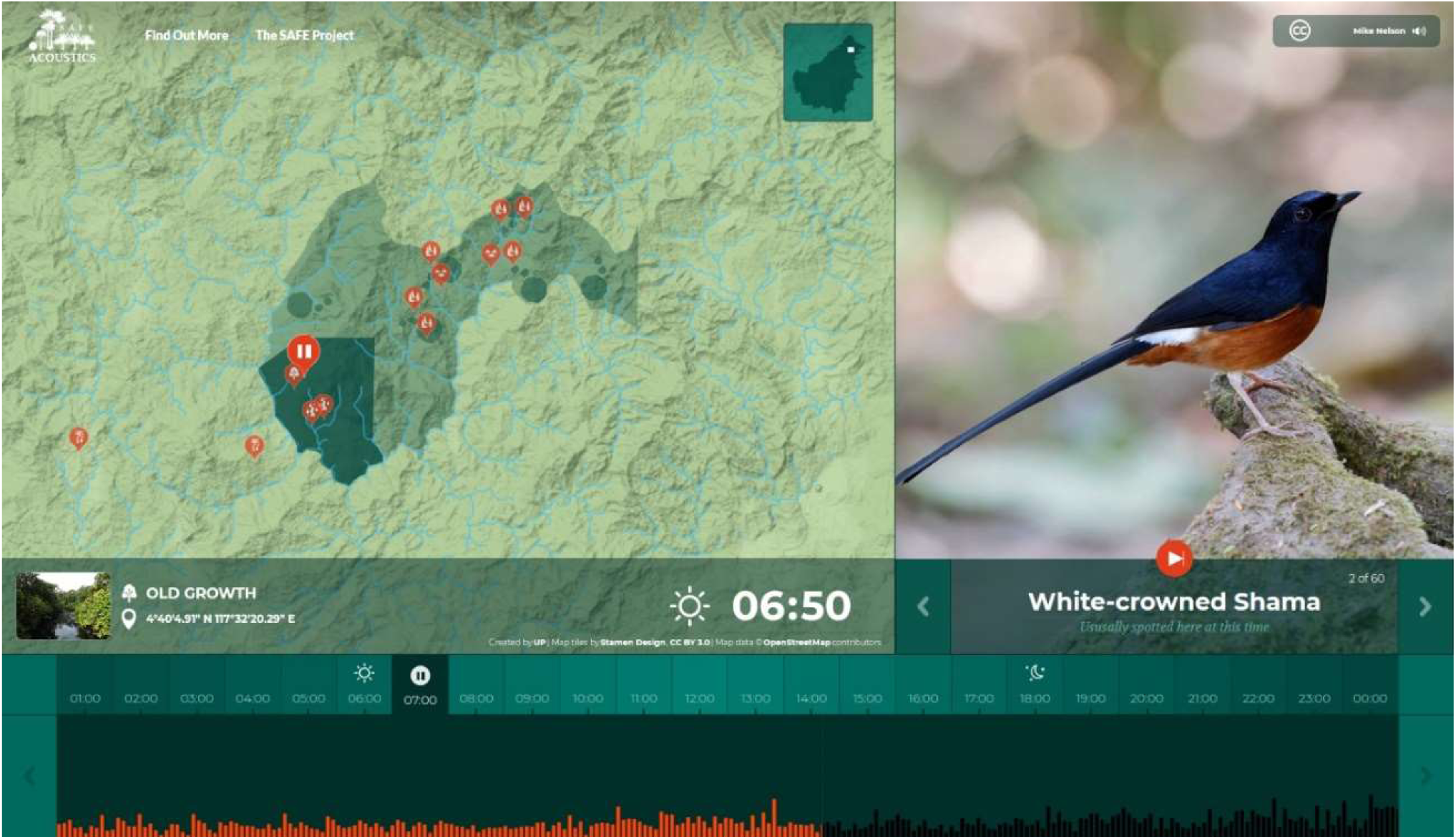
A public-facing website provides a user-friendly interface to engage with audio from our acoustic monitoring network. We developed a ReactJS web app (http://acoustics.safeproject.net) which allows the public to browse a map of our monitoring network (left), listen to audio across the day from each of the recorders (bottom), and explore the species communities present at each location (right).

### Scope for real-time automated analyses and alerts

Whilst the website encourages visitors to make qualitative comparisons of soundscapes, the same infrastructure can be built upon to provide quantitative insights into ecological health^6^. We plan on developing continuous analysis methods which monitor habitat quality, autonomously detect environmental ‘anomalies’ and record the presence of certain species in real-time^6,13^.

An example extension we developed to perform real-time analysis of the incoming audio is as follows. A virtual machine (VM) hosted on Amazon Web Services contacts the acoustics-db API once per day to access the latest audio uploaded from the SAFE Acoustics network. The audio is then processed by an automated audio analysis algorithm on the VM (in this demonstration a speech to text conversion is performed), and the results are published on Twitter via their API (https://twitter.com/BotJungle). A similar approach – employing a more ecologically focussed analysis – could be taken to update a web-based dashboard, or send email alerts to communicate analysis results to land managers or scientists in real-time.

## Conclusion

Over 16,500 hours of audio (49,468 files) have been uploaded from the SAFE Acoustics monitoring network, and the public-facing website has hosted over 13,000 unique visitors. Each stage of the pipeline, from autonomous monitoring hardware to the website user-interface, has been made fully open-source, along with detailed documentation available online. We believe that our scalable, robust design will provide an exemplar of a successful ecological monitoring network for other scientists to build upon. Our work will ultimately accelerate the rate at which wide-scale, fully autonomous monitoring moves from a long-held aspiration to a genuinely impactful tool in ecology.

## Acknowledgements

We thank the research assistants and staff at the SAFE project camp, in particular Ryan Gray, Jani Sleutel, Adi Shabrani, Nursyamin Zulkifli, Unding Jami, and the canopy access team. We thank Henry Bernard for his collaboration. We acknowledge the support of the Imperial College Advanced Hackspace. This project was supported by funding from WWF UK (Biome Health Project), the Sime Darby Foundation, the Engineering and Physical Sciences Research Council [EP/R511547/1], and the Natural Environment Research Council [NE/K007270/1]. S.S.S. is also supported by Natural Environment Research Council through the Science and Solutions for a Changing Planet DTP.

## Author’s contributions

S.S.S., R.M.E., N.S.J., A.S., L.P., and C.D.L.O. contributed to the conceptualisation and final implementation of the study. C.D.L.O., A.S., and S.S.S. developed the open source software for the monitoring hardware, acoustics-db server and SAFE Acoustics website. S.S.S. and R.M.E. led the field deployments of the monitoring devices. S.S.S. led the manuscript writing process with revisions provided by all authors.

